# The Most Informative Neural Code Accounts For Population Heterogeneity

**DOI:** 10.1101/112037

**Authors:** Elizabeth Zavitz, Nicholas SC Price

## Abstract

Perception is produced by ‘reading out’ the representation of a sensory stimulus contained in the firing rates of a population of neurons. To examine experimentally how populations code information, a common approach is to decode a linearly-weighted sum of the neurons’ firing rates. This approach is popular because of its biological validity: weights in a computational decoder are analogous to synaptic strengths. For neurons recorded in vivo, weights are highly variable when derived through machine learning methods, but it is unclear what neuronal properties explain this variability, and how the variability affects decoding performance. To address this, we recorded from neurons in the middle temporal area (MT) of anesthetized marmosets (*Callithrix jacchus*) viewing stimuli comprising a sheet of dots that moved coherently in one of twelve different directions. We found that high gain and direction selectivity both predicted that a neuron would be weighted more highly in an optimised decoding model. Although learned weights differed markedly from weights chosen according to a *priori* rules based on a neuron’s tuning profile, decoding performance was only marginally better for the learned weights. In the models with a *priori* rules, selectivity is the best predictor of weighting, and defining weights according to a neuron’s preferred direction and selectivity improves decoding performance to very near the maximum level possible, as defined by the learned weights.

**New & Noteworthy:** We examined which aspects of a neuron’s tuning account for its contribution to sensory coding. Strongly direction-selective neurons were weighted most highly by machine learning algorithms trained to discriminate motion direction. Models with *a priori* defined decoding weights demonstrate that the learned weighting scheme causally improved direction representation by a neuronal population. Optimising decoders (using machine learning) lead to only marginally better performance than decoders based purely on a neuron’s preferred direction and selectivity.

## INTRODUCTION

Perception reflects the interpretation of the aggregate activity across a population of neurons. Firing rates from ensembles of hundreds of neurons must be combined to accurately represent stimulus properties. We understand some aspects of how a perceptual ‘readout’ may be produced from this population activity, such as how the responses of neurons with different tuning preferences can be weighted and evaluated in order to make discriminations (Seung and Sompolinsky 1993; Graf et al. 2011; Shamir 2014) or to identify where a stimulus lies on a continuous scale (Georgopoulos et al. 1986; Salinas and Abbott 1994). Often, the neural population is idealized using identical tuning curves to uniformly tile the stimulus space and it is reasonable for every neuron to contribute equally to the readout. However, cortical neurons are diverse: within a single brain area they vary in sharpness of tuning, selectivity, overall firing rates, and trial-to-trial variability (Ecker et al. 2011). Here we address the problem of optimal coding in a heterogeneous population by asking which neurons are the most informative, whether their usefulness can be predicted by their response properties, and to what degree we can expect their contributions to coding to be optimized.

The firing rate of a given neuron depends on the activity from, and relative weighting of, each input. This weighting is governed by the strength of synaptic connections from each input neuron. In sensory decision making, the magnitude of a neuron’s synaptic weight can be taken to indicate the amount to which it contributes to a decision. A decoder emulating a decision making process represents the activity of readout neurons that receive inputs from a population of sensory neurons, and its weights are analogous to synaptic weights (Jazayeri and Movshon 2006). To decode neural activity computationally, each neuron’s responses may be assigned multiplicative weights *a priori*, based on known attributes of the neurons such as preferred direction. Alternately, weights may be derived with machine learning approaches. As expected (Pouget et al. 2003), the weights that individual neurons are assigned by decoders trained to perform fine discrimination tasks are highest when the individual tuning curves have the most discriminatory power with respect to the tuning preference of the neuron (Graf et al. 2011; Berens et al. 2012). Even after accounting for this relationship, what factors account for the considerable amount of variability remaining in the weights that neurons in heterogeneous populations are assigned?

Individual neurons in the primate middle temporal area (MT) are tuned for motion direction; each neuron has a preferred direction, which evokes the highest average spiking rate, and average spiking shows an approximately Gaussian dependence on stimulus direction (Albright et al. 1984; Petersen et al. 1985; Newsome and Pare 1988; Lui et al. 2007). Because of this individual tuning, the ensemble of responses from neurons in MT produces a robust representation of the direction of visual motion (Georgopoulos et al. 1986; Bialek et al. 1991; Jazayeri and Movshon 2006; Graf et al. 2011; Chen et al. 2015; Goddard et al. 2017). In this paper, we used the responses from populations recorded in area MT of the marmoset visual cortex to examine how the direction of moving patterns is encoded by neural populations. We used weights optimized through machine learning to determine which neurons are most informative about stimulus direction, and compared these weights to the tuning properties of each neuron in order to predict which neurons are informative. We found that direction selectivity predicted the learned weights of the recorded neurons, and could be used to enhance the performance of decoding models in which weights are predefined, rather than learned. By adopting a modeling approach that allows us to explore causal contributions of applying weights according to a neuron’s tuning parameters, our results show that weighting those neurons that are most selective is the best way to improve decoding performance in both fine and coarse discrimination tasks.

## MATERIALS AND METHODS

### Electrophysiology

We performed extracellular recordings in four anaesthetized, adult New World monkeys (3 male, 1 female; common marmoset, Callithrix jacchus). This data set and recording methods have been previously described in detail (Zavitz et al. 2017). All experiments were conducted in accordance with the Australian Code of Practice for the Care and Use of Animals for Scientific Purposes and were approved by the Monash University Animal Ethics Experimentation Committee.

Briefly, anesthesia was induced with alfaxalone (Alfaxan, 8 mg/kg), and a tracheotomy and vein cannulation were performed. The animal was artificially ventilated with a gaseous mixture of nitrous oxide and oxygen (7:3) and infused intravenously with a maintenance solution including opiate anesthetic (sufentanil) and paralytic (pancuronium bromide). The eyes were treated with atropine and phenylephrine hydrochloride eye drops, and corneas protected with a contact lens selected to bring a display at 350 mm into focus. We performed a craniotomy and durotomy over the contralateral middle temporal area (MT) and implanted a 10×10, 96 channel “Utah” array (Blackrock Microsystems) with 1.5 mm electrodes to a depth of 1 mm. The implantation location was selected using gross anatomical landmarks, and verified based on recorded receptive field characteristics and post-mortem histology. We recorded neuronal activity using a Cerebus system (Blackrock Microsystems) with a sampling rate of 30 kHz. The raw voltage signal was high-pass filtered at 750 Hz, and multiunit activity was detected online based on threshold crossings. Our analyses are based on offline spike-sorted multiunit activity from visually responsive channels in four animals, where a unit was visually responsive if its firing rate during the 500 ms stimulus period at the stimulus direction that evoked the highest response was significantly different from its firing rate during the preceding 500 ms blank as determined by a t-test with a criterion of 0.05. We trial-shuffled responses from all units in each animal to remove any trial-by-trial correlations in cross-channel spiking activity, and then combined all units to produce a single (n = 291) pool from which we could draw smaller hypothetical neuronal populations. Although we have shown that spike count correlations between pairs of neurons impact coding (Zavitz et al. 2017), considering the trial-by-trial correlations within a population is not relevant to the analyses here, which focus entirely on the role of trial-averaged tuning properties. For this reason, shuffling and combining recorded data across cases improves the statistical power of our analyses without impacting on the research question at hand. We previously analysed single and multiunit tuning properties separately with respect to a twelve-alternative classification problem, and found no qualitative difference between multi and single units (Zavitz et al. 2017).

### Visual stimulation

We presented stimuli on a VIEWPixx 3D (1920×1080 pixels; 520×295 mm; VPixx Technologies) positioned at a viewing distance of 350 mm. The stimulation protocol has been previously described (Zavitz et al. 2017). Briefly, we displayed a sheet of white dots on a black background that moved coherently, with no noise, in 1 of 12 equally-spaced directions and 1 of 3 speeds (5, 10, and 20 degrees per second). The stimulus moved in one direction for 500 ms and was followed by a black screen for 500 ms before another, random, direction was presented. Each direction was presented 120 times. We used Psychophysics Toolbox to generate stimuli in Matlab (Brainard 1997; Pelli 1997; Kleiner et al. 2007).

### Characterizing units

We characterized our recorded neurons by their preferred direction, bandwidth, Fano factor, gain, and direction selectivity. A raw tuning curve was constructed based on the mean response to each stimulus direction. Gain was calculated as difference between the maximum and minimum responses. Preferred direction was calculated as the vector sum of the tuning curve, and direction selectivity as its vector magnitude (Equation 1). The bandwidth was computed based on the full width at half height of a Von Mises function fit to the tuning curve. Fano factor was calculated as the average of the spike count variance for each direction divided by its mean.

To calculate a direction selectivity index, we used the following formula:

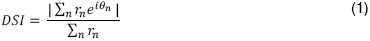

Where r_n_ is the mean response evoked by stimulus direction θ_n_. This measure of direction selectivity depends on the relationship between gain, bandwidth, and minimum firing rate, so it is not independent of our measure of gain. Using both gain and selectivity allows us to compare the differential effects of rate and a more complete description of tuning curve shape (DSI). That these measures are related is not a problem for subsequent analyses, as our goal is to best predict which weights should be applied to each neuron, and to determine whether assigning weights in this way can produce near to optimal decoding.

### Linear discriminant decoding

We used a binomial logistic regression model to decode neuronal responses. The stimulus direction was decoded at multiple time points throughout the stimulus period based on spike counts measured in time windows of 4-128 ms and with a range of offsets relative to stimulus onset. The spike counts for each neuron were weighted, then integrated across neurons and nonlinearly transformed (Figure 1A). This output was compared to a criterion to determine the direction reported by the model. For optimization of weights, we used the glmnet statistical software package in Matlab (Friedman et al. 2010; Qian et al. 2013), configured to use a ridge regression for regularization. The models were trained on 80% of the stimulus trials by refining weights to optimize performance via penalized maximum likelihood. The performance we report is the decoding performance on the remaining 20% of trials. The training trials were randomly selected each time a decoder was trained. In different experiments we varied the sizes of the neuronal populations and the size of the spike integration window, but unless otherwise specified we used subpopulations of 20 neurons, and integrated spikes for 30 ms.

**Figure 1.**
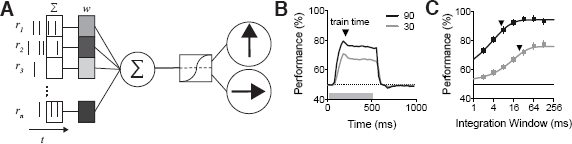
**(A)** Diagram of the linear decoding scheme used here. Spike counts within specific time windows (r_i_) from n neurons are weighted, summed across neurons, nonlinearly transformed and compared to a decision threshold. The weights are adjusted to optimize performance when the model is trained on a subset of 80% of the available trials and can then be tested on the remaining trials, at the same or a different time point. **(B)** The population representation of motion, as measured by linear decoding performance, emerges quickly after stimulus onset at time = 0, and is sustained throughout the 500 ms of stimulus presentation (gray bar). For further analyses with recorded neural data, we selected a single time point in the stimulus response (200 ms). **(C)** Decoding performance reaches ceiling with a sufficiently large spike counting window. Mean performance from 100 resampled subpopulations of 20 recorded neurons (points, error bars indicate 95% CI) is fit with a Weibull function to interpolate an ideal window. For both decoding problems, we used a small enough window to ensure that the weights were optimized (black triangles).

To perform two-alternative discriminations on a data set collected using 12 stimulus directions, we separately decoded only the trials relevant to a specific pairing of directions. For example, in the coarse two-alternative discrimination, we trained models to decode 0° vs 90°, 30° vs 120°, etc. Every direction pairing was decoded for every subpopulation, and decoding results were combined across pairings.

### Fitting and using a priori decoding models

To create a priori decoding models, we used the same procedure described above, except instead of learning the optimal weights for each neuron and class, we applied the weights predicted by the Gabor weighting model in Equation 2:

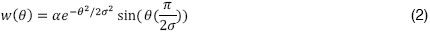

We used a Gabor weighting to perform direction discrimination. The relevant weight is determined by three parameters: the gain (α); the spread (σ) of the Gabor; and the difference between a neuron’s preferred direction and the discrimination boundary between the two classes. The zero-crossing of the Gabor was fixed at θ_pref_- θ_decisionBound_ = 0. The frequency of the sine wave was set as a ratio of the Gaussian’s standard deviation, so that one full cycle fit within the Gaussian envelope.

### Simulation

Populations of neurons were simulated based on the distributions of gain, bandwidth, Fano factor, and spontaneous activity that we observed in our recorded populations. A Gaussian tuning curve was produced with the specified gain, bandwidth and spontaneous level, and spike counts were generated for a given trial using a negative binomial distribution with parameters determined by each neuron’s tuning curve and Fano factor. These spike counts were then decoded in the same manner as our recorded populations of neurons.

## RESULTS

We recorded spiking activity from neurons in area MT of the marmoset while the animal viewed a stimulus that displayed a field of dots moving in one of 12 motion directions for 500 ms, followed by 500 ms of a blank screen. We assessed the population representation of motion direction by applying a linear discriminant decoding analysis, which uses spiking information from multiple neurons to predict the direction of motion that was displayed. We recently demonstrated that some neuronal tuning properties are correlated with the weights assigned to these neurons in a multinomial (12-alternative forced choice) linear decoding scheme (Zavitz et al. 2017). Here, we extend those findings to two binomial linear decoding schemes performing either fine (30° separation) or coarse (90° separation) discrimination (Figure 1A), and test whether the correlations we observed lead to a causal improvement in direction coding.

### Decoding weights depend on the properties of individual neurons

To determine which neurons contribute the most to the neural code, we began by assessing how neurons recorded from marmoset MT are weighted in a linear discriminant analysis, and then correlated those weights against several properties of the tuning curve of each neuron. This procedure allowed us to determine how weights in a decoding scheme relate to each cell’s response properties.

#### Factors influencing decoding performance

Populations of MT neurons contain a strong representation of visual motion direction, and if spikes from many neurons (r_1_… r_n_) are integrated for an extended period of time (t), predictions of the direction of motion based on neuronal activity quickly reach ceiling as the number of neurons or time are increased. To examine aspects of the quality of the population code, it is important for performance to be below ceiling so that we may detect variations in performance following experimental manipulations, and so that the weights reflect a model that is optimized for the given population, as opposed to simply sufficient for ceiling performance. To this end, we present results for a small subpopulation of neurons (n = 20), with spiking activity integrated over a window that produces sub-ceiling performance.

We find robust direction coding (Figure 1B), with above chance performance on fine (30°, gray) and coarse (90°, black) discrimination by 100 ms after stimulus onset. In this instance, we trained the model on spikes counted 200 ms after stimulus onset, but we have shown previously that the direction representation in MT is quite stable over time (Zavitz et al. 2016, 2017). The performance of the decoder improves as spikes are counted over increasingly large windows (Figure 1C), as measured for 100 resampled subpopulations of 20 neurons each. We characterized this relationship by fitting a cumulative Weibull distribution to the performance as a function of integration window (solid lines), and determined which integration window produced 90% of ceiling performance for the average sub-population of 20 neurons. These windows, to the nearest millisecond, are indicated by black arrows: 23 ms for a fine discrimination, and 6 ms for a coarse discrimination.

These results serve to show the context for our analyses of neuronal recordings. Further analyses are performed on weights obtained when training and testing the models 200 ms after stimulus onset, using the illustrated integration window.

#### Properties that influence weights

The performance of a decoder on each trial is determined by the interaction between each neuron’s response and its assigned weight. A neuron’s weight reflects its contribution to the population representation, so neurons that are highly weighted have a larger influence over coding, and perception (Jazayeri and Movshon 2007). To determine which tuning properties predict a large contribution, we first examined weight relative to the proximity of the preferred direction to the decision boundary (i.e. in a 0° vs 30° discrimination, the decision boundary is 15°). We show that weights are highest for neurons with preferred directions approximately 90° from the discrimination boundary, and lowest for neurons with preferred directions at the discrimination boundary (Figure 2A). This is consistent with previous observations in V1, where weights in an orientation discrimination task are highest for cells with preferred orientations near 45° from the discrimination boundary in a 90° orientation discrimination task (Graf et al. 2011), given that direction decoding happens in a 360° space, while orientation decoding happens in a 180° space.

**Figure 2.**
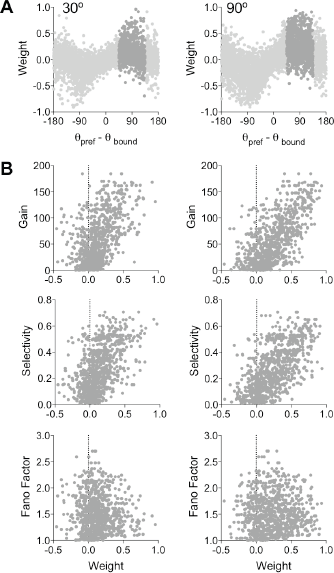
**(A)** Weights are plotted against the difference between the neuron’s preferred direction and the decision boundary. On average, the weights that neurons are assigned show a systematic relationship with this direction difference, but also show a high degree of variability. Shaded values indicate the values that contributed to the subsequent correlation analysis: those whose preferred direction was within 45° of the peak value. Each neuron is depicted 12 times because each neuron is weighted in 12 different discriminations (0° v 30°, 30° v 60°, etc.). **(B)** Correlation between decoding weight and gain, selectivity or Fano factor. Only neurons with dark shading in (A) are considered. Neurons with high gain or high direction selectivity tend to have high weights. In fine discriminations, neurons with high variability tend to have somewhat lower weights. Each neuron is depicted at least twice in these plots, and a neuron whose preferred direction is 90°±5° from the decision boundary is depicted three times.

The relationship between weight and preferred direction is nonlinear, so to permit us to examine how weight was affected by other factors, we selected only those weights that were learned when a neuron was most likely to be highly weighted: when its preferred direction was within 45-135° of the discrimination boundary (dark points, Figure 2A). After selecting those weights, we measured partial Spearman correlations between weight and a subset of tuning properties: gain (height of tuning curve relative to spontaneous rate in spikes per second), selectivity (direction selectivity index), and variability (Fano factor). We found that gain and selectivity were positively correlated with weight in both fine (r_gain_ = 0.30, p = 2.9×10^−19^; r_DSI_ = 0.28, p = 7.3×10^-17^) and coarse (r_gain_ = 0.41, p = 5.2×10^−36^; r_DSI_ = 0.37; p = 5.6×10^−30^) discriminations. Fano factor was found to have a significant negative correlation with weight in the fine discrimination (r_FF_ = −0.07; p = 0.02), but not in the coarse discrimination (r_FF_ = −0.01, p = 0.78). This suggests that neurons with responses that are larger and more selective are weighted most strongly. It also suggests that neurons with lower variability (and thus lower Fano factors) are assigned somewhat higher weights. This is consistent with our previous observations in a multi-class identification paradigm (Zavitz et al. 2017).

These findings are in line with our previous findings, and with our expectations as neurons with the highest discriminatory power should contribute the most to sensory discriminations. The description above, derived from experimental data, does have some important caveats. First, it is based on a large, but limited, population of cells. To describe general principles, we require a more general population. Second, our description is not causal. We have shown that certain properties are associated with higher weights in a machine learning model, but we have not yet shown that assigning higher weights to neurons with these properties causally improves discrimination performance.

### Models fit to simulated neurons describe weighting functions by tuning property

Our finding that some response properties correlated with weight in a linear decoder was based on a relatively small number (291) of recorded neurons. To expand the population of neurons from which we are drawing inferences, we simulated 500 populations of 20 neurons each, whose tuning properties were drawn from the distributions of those we recorded. This approach allowed us to define a weighting function that describes the relationship between the decision and the preferred direction, and to systematically determine how the shape of the weighting function is different for neurons with different properties. We did this in two stages: first, we used a homogeneous population to fit a model that describes the relationship between preferred direction and weight; then we simulated heterogeneous populations and examined how the parameters of the model depended on the tuning properties of the neurons. As shown in this paper and previously, for a given relationship between the preferred direction and the class or decision boundary, weights in heterogeneous populations are variable (e.g. Figure 2A). This variability precludes us from fitting a model that describes a straightforward relationship between the preferred direction of the neuron, the class or decision boundary, and the weight the neuron is assigned in a decoder. To overcome this, we first simulated a homogeneous population, in which each neuron has a different preferred direction, but the same tuning curve shape. Note that only gain and Fano factor will be necessarily uncorrelated. The selectivity that emerges from a tuning curve with a given gain, bandwidth, and spontaneous rate is, in practice, still correlated with gain (r = 0.71, p = 1.0×10^-77^).

We measured the distributions of the properties of our recorded population in order to simulate responses from a population with analogous statistics. We quantified gain, spontaneous rate, bandwidth, and Fano factor (Figure 3A). Gain, spontaneous rate, and bandwidth together determine the direction selectivity that is ultimately observed. To simulate a homogeneous population, every unit had the same tuning curve shape, as described by the medians of the populations distributions (Figure 3A), and preferred directions were chosen to uniformly tile 360°. We used simulated responses from 100 homogeneous populations of 20 neurons over 400 trials each with the same decoding approach as previously. This produced a series of learned weights for a fine (Figure 4A) and coarse (Figure 4B) discrimination. Because the firing rate on a finite number of trials is drawn from a negative-binomial distribution (whose mean-to-variability ratio is determined by the Fano factor), the weights are still variable, but they are well-described by a fitted Gabor function with two free parameters: amplitude and spread (r^2^=0.92 for the fine discrimination, and r^2^ = 0.98 for the coarse discrimination) (Eq. 2; Figure 4A & B, lines).

**Figure 3.**
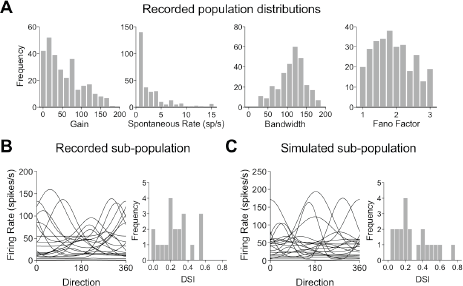
**(A)** Distribution of the tuning properties (gain, spontaneous rate, bandwidth, and Fano factor) for 291 recorded neurons, which were used to constrain the properties of simulated neuronal responses. **(B)** Tuning curves and direction selectivity indices for a subpopulation of 20 recorded neurons. **(C)** Example of tuning curves generated by the model for a population of 20 neurons, and the distribution of their measured DSIs. Note that the observed distribution of DSI emerges from basing the simulated populations on the tuning properties in (A) only.

**Figure 4.**
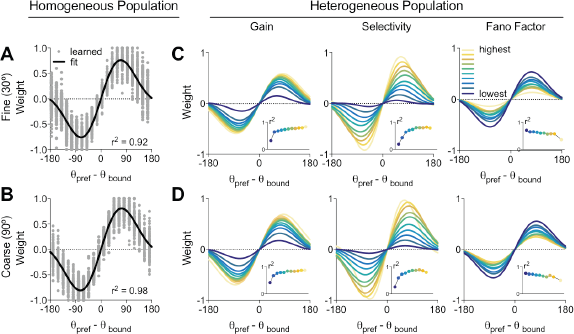
**(A)** weights learned for responses in homogeneous synthetic populations (dots) in a fine discrimination tasks are well-described by a fitted Gabor function with two parameters: amplitude and spread. **(B)** As in A for a coarse discrimination. **(C & D)** Fits of a two-parameter Gabor to weights of heterogeneous subpopulations of simulated neurons. Fits have been made to subsets of neurons binned based on different tuning properties: gain, selectivity and Fano Factor. The dark blue functions are fitted to weights for neurons with the lowest of this property, the yellow functions to the highest. Inset panels show r^2^ for each model.

We then simulated responses from 500 heterogeneous populations of 20 neurons whose properties were drawn randomly with replacement, from the full distribution of recorded properties (Figure 3A), with preferred directions selected randomly from a uniform distribution. These simulated populations are comparable to the recorded subpopulations, and produce similar distributions of selectivity (Figure 3B, C). As before, we used a generalized linear model to learn decoding weights for each subpopulation, then partitioned the weights for the 10,000 simulated neurons into bins based on their gain, selectivity, and Fano factor. For each of these three features, we separately partitioned neurons into 11 evenly spaced bins, and fit a Gabor model to relate the weights to a neuron’s preferred direction for each bin and metric (Figure 4C & D). Following this approach, rather than a single weight model for the population (as in Figure 4A & B), there are 11 distinct models, which capture some of the variability in weights across a population (Figure 4C & D, insets). Model fit is good enough to be used to generate predicted weights in all cases (r^2^ values range from 0.20 for bins containing cells with very low selectivity, to 0.88 for bins containing cells with the highest selectivity). The models fit to the neurons with the lowest gain and selectivity, and the highest Fano factor fit less well than the others. This is likely because those cells with the weakest tuning and most variable responses do not follow any particular weighting function with respect to preferred direction.

The results of this analysis are consistent with those derived from the correlation analysis carried out previously (Figure 2), but are much clearer because they are based on a larger underlying pool of response characteristics (simulated neurons). Although gain was strongly correlated with weight, functions fitted to neurons with the highest gains (Figure 4C & D, leftmost column, yellow) are not very different from those fitted to those with moderate gains (green). We can see that the best fitting function depends strongly on selectivity, across the entire range of neurons. Furthermore, it is apparent that the most selective cells are not just better-fit by higher amplitude functions, these functions also weight neurons more highly closer to the discrimination boundary, particularly in the fine discrimination task, as is ideal in a fine discrimination (Jazayeri and Movshon 2006; Graf et al. 2011). This is likely because the bandwidth of a highly selective neuron is narrow enough to be optimized in this way. It is also telling that although the correlation between Fano factor and weight was weak (and not statistically significant in the coarse discrimination, Figure 2B), the weighting functions have a small but very systematic reliance on Fano factor.

The weight values depicted in Figure 4C & D are appropriate for neurons sorted into coarse bins by their tuning properties, but ideally we would like to tailor a decoding weight according to the individual properties of an arbitrary neuron. This is possible because the amplitude and spread of each model change systematically as a function of the tuning properties at hand (Figure 5). This means that it is possible to predict the shape of the best fitting weight function based only on known tuning properties. In order to interpolate model parameters between the bins we used for the previous analysis, we fit the Gabor model parameters (amplitude a and spread σ in Eq. 2) as a function of tuning property (gain, selectivity, Fano factor) with second order polynomials (Figure 5).

**Figure 5.**
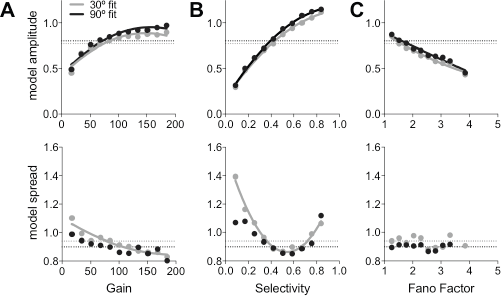
**(A)** Values of the model amplitude α and spread σ parameters (Eq. 2) for simulated neurons with the specified gain (points) in a fine discrimination (30°, gray) and a coarse discrimination (90°, black). These are the parameters that generate the functions depicted in Figure 4B. The value of each parameter is systematically related to the tuning properties of the neurons, which is well captured with a second-order polynomial fit (line). Those fits shown with solid lines are those that were used to determine the parameters of the Gabor model. Dotted horizontal lines show the median values of either amplitude or fit for all fine and coarse models. These medians were used when the Gabor model fit did not meet our inclusion criteria. **(B)** As in (A) for neurons grouped by direction selectivity. **(C)** As in (A) for Fano factor. The parameters, and polynomial fits, are qualitatively consistent between the fine and coarse discriminations.

We used these polynomial fits to predict the amplitude and spread of the model weights if (1) they were better than a criterion of r^2^ = 0.9, and (2) they passed a shuffle correction test (α = 0.01) requiring that the fit was significantly better than a second order polynomial fit to shuffled data. These criteria ensure that the fitted polynomial accurately describes the shape of the Gabor model parameters and that the shape of the parameters depends systematically on the property under consideration. In further analyses, parameters whose polynomial fits did not pass both criteria were fixed at the median value for all tuning properties (dotted lines in Figure 5).

All of the tuning properties we examined significantly and systematically modulated the amplitude of the Gabor model for decoding weights in both the fine and coarse discriminations. Only gain and selectivity modulated the spread of the function, and, as expected, this was the case only for the fine discrimination. As is qualitatively evident in Figure 4, the fit parameters are most strongly modulated by the selectivity of the cell.

These results show that larger weights are learned for neurons with higher gain, greater selectivity and lower variability, and outline quantitatively what weight is expected for a given preferred direction, selectivity and Fano factor. This quantitative description allowed us to set weights for our recorded population of neurons based on these properties alone instead of using a supervised learning algorithm.

### Weighting neurons according to their tuning properties causally improves decoding performance

In the previous section, we simulated populations of neurons in order to produce models that describe how the weights derived using machine learning algorithms relate to the tuning properties of arbitrary individual neurons. A limitation of this approach is that we were only able to correlate the tuning properties of recorded neurons and their weights. Therefore, in this section, we will use these models to address whether assigning higher weights to neurons according to their tuning properties causally improves discrimination performance.

To examine the direct impact of the relationship between tuning properties and weights on performance, we define weights for the neurons we recorded from marmoset MT based on the Gabor functions outlined in the previous section: the parameters for the Gabor function describing the weights (Figure 4) are specified by the polynomial fits to each parameter (Figure 5). This allows us to set a weight for each neuron that depended only its specific tuning properties (Figure 6). The preferred-direction only model uses the median amplitude and spread (as indicated by dashed lines in (Figure 5).

**Figure 6.**
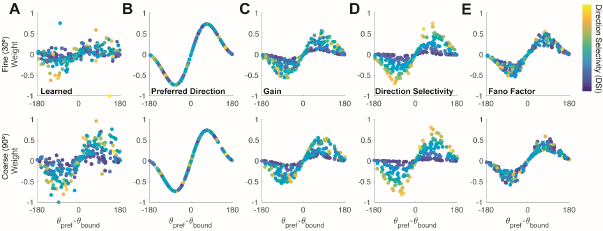
**(A)** Learned weights for one neural population for a fine (top) and coarse (bottom) discrimination. There are 12 weights for each of 20 neurons. **(B)** As in (A), but weighted based on an a *priori* model based only on preferred direction, **(C)** an a *priori* model based on gain and preferred direction, **(D)** an a *priori* model based on selectivity and preferred direction, and **(E)** an a *priori* model based on Fano factor and preferred direction. Note that including preferred direction in the model is necessary, and the preferred direction only model should be considered the baseline, while the learned model should be considered the ceiling. All points are colored according to the direction selectivity of the neuron being weighted.

The learned weights (having been optimized for the chosen population) should produce the best decoding. On the other hand, for direction decoding to happen at all, neurons must be weighted based on their preferred direction, so a model incorporating preferred direction should accurately predict direction, but with the weakest performance. Within this range, we can compare how other models including those that account for gain, selectivity, and Fano factor perform to give insight into which neuronal properties causally affect decoding. We trained decoders on the responses of 100 subpopulations of 20 neurons each, sampled with replacement from our 291 recorded neurons. For each subpopulation, we compared performance with the learned weights (Figure 6A) to performance with weights set according to four different models. We were surprised that the optimal learned weights display remarkably little structure. The preferred direction only model (Figure 6B) is the model typically considered in population decoding. Regardless of any other properties, neurons are weighted based on the relationship between their preferred direction and the class being decoded. To determine the potential benefit of accounting for additional response properties, we modified the weighting function for each neuron based on the polynomial fits for each parameter (Figure 5).

We used these weighting functions to decode the stimulus direction at a single time point (200 ms after motion onset), as before, with each of the models in Figure 6. The performance in the fine discrimination ranges from 64.6% (SD 8.9%) with the learned weights to 62.3% (SD 8.6%) with weights set according to the preferred direction only (Figure 7A). Performance in the coarse discrimination ranges from 74.8% (SD 8.5%) for the learned model weights to 71.2% (SD 9.1%) for the Fano factor model (Figure 7B). To examine the improvement that each model offers, we compared the distributions of performances to the model using the learned weights, Figure 6A. The performance distributions may be paired because we used different weights on exactly the same subpopulations and test trials. Thus, we took the difference between each model and the model where the weights were learned, and converted these distributions of differences into a z-score. To test the statistical significance of these differences, we compared models pairwise using a z-test, using a Bonferroni correction to account for multiple comparisons. (Figure 7C & D). A z-test allows us to examine significance independently of the number of samples, which is important for comparisons of this kind where the data are resampled an arbitrarily large number of times.

**Figure 7.**
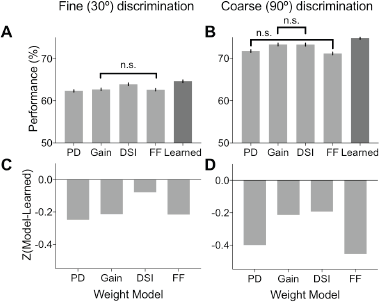
**(A)** Decoding performance in a fine (30°) discrimination for five different weight models. Only pairs of conditions that are not significantly different from one another have been indicated for clarity (i.e. most pairings are significantly different). Error bars show 95% confidence intervals for the mean. **(B)** As in A, for a coarse (90°) discrimination. **(C)** Model performance relative to the decoder that learns optimal weights for each subpopulation for a fine (30°) discrimination. The weights based on preferred direction only decode just over 0.2 standard deviations worse that the learned weighting model. The selectivity (DSI) and variability (FF) models perform significantly better than the preferred direction model, but the gain model does not. The selectivity model performs significantly better than the gain and variability models, and the Fano factor significantly better than gain. **(D)** As in (C) for a coarse (90°) discrimination. In this case, the different models are more comparable, but the model using selectivity is still the best of the tuning-based models, and a model including gain also significantly improves upon the preferred-direction only model. Note that in the coarse discrimination, because none of the models used a spread parameter, the no-amplitude model is the same as the preferred direction model (see Fig. 5, black data points in lower panels).

In both the fine and coarse discriminations, all models performed significantly worse than the theoretical maximum under the learned weights (Coarse discrimination, z-test, p_PD_ = 1.8×10^−137^, p_gain_ = 2.5×10^−26^, p_DSI_ = 3.9×10^−24^, p_FF_ = 2.5×10−^127^; fine discrimination, z-test p_PD_ = 3.9×10^−42^, p_gain_ = 6.5×10^−31^, p_DSI_ = 1.6×10^−05^, p_FF_ = 5.5×10^−31^). However, the baseline model, based on preferred direction alone, is less than 0.5 standard deviations worse (0.46 in the coarse discrimination, 0.25 in the fine). This demonstrates that the bulk of the decoding performance is achieved through simply weighting the neurons appropriately for their preferred direction.

In the fine discrimination (Figure 7A & C), the gain model and Fano factor models are significantly better than the preferred direction model (p_gain_ = 1.6×10^−6^, p_FF_ = 3.0×10^−5^), but not significantly different from each other (p = 0.45). The selectivity model is significantly better than all the others (p_PD_ = 2.2×10^−33^, p_gain_ = 6.3×10^−21^, p_FF_ = 3.1×10^−22^), and produces performance 0.07 standard deviations, (or 0.7 percent correct) worse than the learned weights (1.7-1.3 percent correct better than the other models). This suggests that, in the fine discrimination, the selectivity model captures almost all of the variability in weights essential to performance.

In the coarse discrimination, the Fano factor model is not significantly better than the preferred direction model alone (p = 0.18) (Figure 7B & D). The gain and selectivity models are substantially, and significantly better than the preferred direction model (p_gain_ = 4.7×10^−136^, p_DSI_ = 1.2×10^−64^), although they do not perform significantly differently from one another (p = 0.34).

That the direction selectivity model provides the largest benefit in the fine discrimination condition does not seem surprising: it is the model that can take the best advantage of weighting narrowly tuned cells highly when their preferred directions are close to the discrimination boundary. To examine this in more detail, we evaluated performance on the same data but using only one of the model parameters: either amplitude or spread. The parameter not used was fixed to the median of the measured distribution (Figure 5, dotted lines). This manipulation allows us to test the differential contribution of each parameter. Note that this analysis was only carried out in the fine discrimination condition for models that weight by gain and direction selectivity, because when the spread parameter is not varied, all results are necessarily due to the variation in weighting amplitude. In both the gain and selectivity models, varying spread while keeping amplitude fixed significantly reduces performance relative to the full model (p_gain_ = 1.0×10^−6^, p_DSI_ = 1.3×10^−27^), but varying amplitude while keeping spread fixed does not significantly affect decoding performance (p_gain_ = 0.88, p_DSI_ = 0.24). This suggests that the greater performance in the case of the DSI model is due solely to the wider range of amplitudes it uses.

These results show that accounting for direction selectivity by weighting more selective neurons more highly causally improves the representation of motion in a population code beyond what is attainable by accounting for preferred direction alone. Moreover, this improvement nearly reaches the performance level attained by weights optimized by machine learning in a fine discrimination task. Weighting neurons by gain is a strategy that performs equally well in coarse discriminations.

## DISCUSSION

Taken together, this work has shown that recorded neurons with higher gain and direction selectivity tend to be assigned higher weights for discrimination tasks by machine learning algorithms. Based on simulated, heterogeneous neuronal populations we developed models that can assign weights to neurons based on their tuning properties alone. We then used those models in populations of recorded neurons to show that weights assigned by preferred direction, and to a lesser extent, gain, causally improve the quality of a population code - nearly to the level that machine learning approaches attain. Furthermore, we show that weighting neurons across a large range of amplitudes (rather than carefully refining the spread of the weighing function) is critical to attaining near-optimal decoding performance.

### Discrimination difficulty

In this paper we looked at two discrimination difficulty levels: 30° differences and 90° differences, which we refer to as fine and coarse, respectively. In the total scheme of two-alternative discrimination, both these comparisons are in the middle range, being neither incredibly fine, nor extremely coarse. A discrimination so fine as to be perceptually difficult for a high-contrast grating or high-coherence dot stimulus is on the order of a 3° direction difference (Purushothaman and Bradley 2005). We expect that finer discriminations would generally adhere to the same decoding principles explored here. It is also possible that if we had required the population to make finer discriminations, the narrowness of the weighting function (σ) would be a more important parameter in producing good decoding performance. It is also possible that the spread parameter would improve performance more were it able to vary more widely. Coarser discriminations (i.e. 180° difference in direction) are not well described by this decoding profile, probably because there is less need to integrate information across neurons with overlapping tuning curves (Jazayeri and Movshon 2007). Previous work in this field demonstrated a significant shift in the steepness of the decoding profile (Graf et al. 2011), although they did not examine whether this characteristic of the weights was causally tied to decoding performance. Any inconsistency could be due to the broader bandwidth of neurons we recorded in MT (they used V1 recordings), or because our 30° discrimination was not sufficiently fine.

### Variability in learned weights

We set out to examine factors that explained some of the variability in the weights learned for populations of neurons. Even accounting for these factors, we saw that a substantial amount of variation remained. This could be because we trained the models on a limited number of trials, relative to the inter-trial variability. We used small populations of neurons in this study, small enough that not every population covered the space of stimulus direction equally well. Some of the variability in weights must therefore arise because the usefulness of a neuron in a population code depends on how large a gap it fills within the population.

### Metrics to describe neuronal responses

We selected the measures examined here (gain, selectivity, variability), because they are closely related to the shape of the measured tuning curves (gain, selectivity), or describe the variability of the neuron relative to the mean rate (Fano factor). Gain is not directly analogous to peak firing rate, as it subtracts out the response to the anti-preferred direction. The direction selectivity index (DSI) also accounts for responses to the anti-preferred direction, along with the height and spread of the curve. It doesn’t matter that these metrics are not independent, because we did not seek to isolate a particular property, but to best predict which weights should be applied to each neuron, and whether assigning weights in this way could improve decoding performance to a level near using learned weights.

Prior to large-scale population recordings such as those used here becoming feasible, changes in tuning bandwidth were often taken to indicate improvements in sensory coding. However, the notion of a sharpened tuning curve alone (with gain held constant) does not necessarily improve coding in the context of a larger population (Pouget et al. 1999; Zhang and Sejnowski 1999; Seriès et al. 2004). We chose not to examine bandwidth in isolation, as preliminary results were noisy, and a cell’s bandwidth is incorporated into our measure of selectivity. It has recently been demonstrated that the precise number of spikes in a fixed interval (rate) is what determines how and when a neuron is informative, and that selectivity does not matter (Sun and Barbour 2017). These results are not necessarily in conflict with our own, as our measure of selectivity (DSI) does depend on number of spikes the neuron fires, we did not explicitly examine rate, and we even removed some aspects of rate through our gain measure (via the floor in the firing rate set by the response to anti-preferred direction motion).

### Role of Spike-count Correlations

Correlated trial-by-trial variability is undoubtedly related to the quality of the population code for any represented quantity (Kohn et al. 2016). In this study, we sought to examine the role of tuning independently of correlated variability, so all the data examined were trial shuffled to remove any recorded correlation structure. Correlations in populations this small will tend to weakly facilitate decoding (Franke et al. 2016; Zylberberg et al. 2016; Leavitt et al. 2017; Zavitz et al. 2017), and we have previously shown, using these recorded data, that the tuning properties of the neurons are related to their weighting functions, even in the context of correlated data (Zavitz et al. 2017). In larger populations, correlated variability is more likely to weakly impair decoding, or at least offset any benefits of adding more neurons to a decoding decision (Lin et al. 2015; Kohn et al. 2016), thus we felt it appropriate to ignore trial-by-trial correlations here.

### Population Size and Optimisation

Given the vast population of neurons in the mammalian cortex, creating a population code that is as accurate as possible could be achieved by pooling across large numbers of sensory neurons. However, our results demonstrate that even a very small number of neurons operating at biologically plausible integration windows are very good at representing motion information (Newsome et al. 1989). Our approach, and others like it, are limited as we employ neural data outside of the context of behaviour. We have assessed the information available to a read out system, and have demonstrated that the most effective way to extract it depends on the tuning properties of the neurons.

However, we still do not know how the information is extracted and employed by a behaving organism in order to make perceptual decisions or initiate visually guided movements.

## AUTHOR CONTRIBUTIONS

EZ ran experiments, developed analysis procedure and simulations, interpreted results, and wrote the paper. NP designed and ran experiments, provided insight into analysis procedure and the interpretation of results, and contributed to the paper.

## ACKNOWLEDGMENTS

We thank Marcello Rosa, Hsin-Hao Yu, Saman Haghgooie, and Amanda Davies for their help running the electrophysiology experiments. We thank Hsin-Hao Yu and Katrina Worthy for histological tissue processing.

We thank Janssen-Cilag Pty Limited for the donation of sufentanil citrate.

